# Single-component optogenetic control enables human portrait formation in cells

**DOI:** 10.1101/2025.11.13.688218

**Authors:** Fatma Yilmaz-Atay, Merve Kocoglu, Busra Nur Cevik, Asli Ceren Eker, Eda Demirozcan-Kurun, Nuri Ozturk

## Abstract

Continued exploration of novel photoreceptors is essential for advancing optogenetic tools, as differences in their reaction mechanisms and spectral sensitivity could enhance the versatility of available systems. While circadian clock photoreceptors represent promising candidates, only *Arabidopsis* Cryptochrome 2 (AtCRY2) has been widely adopted in current applications. Notably, both the light-dependent interaction of AtCRY2 with the CIB1 protein and its homodimerization have been exploited for diverse optogenetic applications. In our search for alternative circadian photoreceptors for optogenetic control, we aimed to develop a system functioning without the co-expression of interacting partners. In this context, *Drosophila* Cryptochrome (DmCRY) was a promising candidate due to its well-characterized photoreceptor function and defined light-dependent reaction mechanism. Upon illumination, DmCRY undergoes a conformational change that enables interactions with Jetlag and Timeless proteins, facilitating circadian clock resetting. DmCRY is also targeted for degradation via ubiquitylation by Jetlag and/or BRWD3, ensuring resetting occurs only once each morning. Notably, DmCRY is susceptible to light-induced degradation even when expressed in mammalian cells. Leveraging this property, we fused the Tet repressor (TetR) to the C-terminus of DmCRY to enable light-dependent regulation of gene expression via TetR-responsive elements. We demonstrate that this system modulates expression of Cas9 for gene knockout, dCas9 for transcriptional regulation, and recombinant proteins. Furthermore, using these tools with a photomasking technique, we generated human portrait images from cultured cells. These findings highlight DmCRY as a versatile optogenetic tool capable of controlling cellular processes with standard visible light, with potential for novel optogenetic platforms.

## Introduction

Precise control of cellular processes can be achieved by manipulating specific molecules or proteins within cells, with gene expression being a common regulatory target. Chemical approaches such as the Tet-ON and Tet-OFF systems modulate gene expression through allosteric mechanisms (1). Other strategies use small molecules to mediate the reassembly of split transcription factors or other components (2). Beyond chemical approaches, photoreceptor proteins have been engineered to control cellular processes in a field known as optogenetics (3), which enables light-dependent regulation of cellular functions, including gene transcription. In these systems, a photoreceptor senses light and activates an effector or actuator that regulates specific processes. Effectors can be designed for tunable, reversible, and bidirectional control, with light serving as a noninvasive external input. Advantages of optogenetics include high spatial resolution, minimal off-target effects, rapid reversibility, and fine-tuned regulation (4). Excellent optogenetic tools based on channelrhodopsins achieve optical control through light-gated ion conductance across membranes, enabling precise modulation of neuronal activity (5). On the other hand, non-transmembrane proteins such as circadian photoreceptors which undergo light-induced conformational changes with long-lasting activity may also offer compact and single-component architectures capable of targeting diverse cellular processes. Tools based on light-induced dimerization of *Arabidopsis thaliana* photoreceptor cryptochrome 2 (AtCRY2) and its light-dependent interaction with its partner CIB1 have enabled bidirectional gene expression control in mammalian cells, either by reconstituting split transcription factors or by blocking transcription factor nuclear localization (6). AtCRY2 belongs to the cryptochrome/photolyase family (CPF), which comprises cryptochromes (CRYs) and photolyases (PLs). CRYs contribute to circadian clock function either as regulators or as photoreceptors, whereas PLs mediate light-driven DNA repair (7). Beyond AtCRY2 and AtCRY1, CRYs from diverse taxa also function as photoreceptors, although their reaction mechanisms vary (8). Among animal CRYs, *Drosophila melanogaster* cryptochrome (DmCRY) has been extensively studied for its role in circadian clock resetting (9, 10). The animal circadian clock operates through a transcription translation feedback loop (TTFL) mechanism (11). In the positive arm of the TTFL, Clock and Cycle (Bmal1 in mammals) activate circadian clock-controlled genes (CCGs), two of which constitute the negative arm of the TTFL. The protein products of the negative arm translocate into the nucleus and inhibit the activity of the positive arm, thereby sustaining oscillations (12, 13). In *Drosophila*, the negative arm comprises *Period* and *Timeless* genes. Light-activated DmCRY triggers a pathway that recruits Jetlag to ubiquitinate Timeless protein, leading to its degradation and relieving inhibition on Clock/Cycle, thus initiating a new oscillation cycle (14).

Even though *Drosophila* and mammalian core clocks are generated by relatively similar TTFL mechanism, the resetting mechanism shows great variation. In *Drosophila*, DmCRY functions as the primary circadian photoreceptor (15), whereas in mammals, Cryptochrome 1 and 2 act in the negative arm of the TTFL in place of Timeless (16–18), and melanopsin in specialized retinal ganglion cells mediates photic entrainment (19). Because mammalian CRYs lack photosensory function (20), DmCRY presents an attractive optogenetic candidate for use in mammalian cells without perturbing endogenous circadian oscillations, analogous to AtCRY2-based approaches. While AtCRY2 and its light-dependent interactions have been exploited in mammalian optogenetics, no functional DmCRY-based system had been developed. Importantly, DmCRY is degraded after signaling to Jetlag and Timeless in *Drosophila* (21, 22) and can also undergo light-induced degradation in mammalian cells independent of its *Drosophila-*specific interactors (23). This property suggested that DmCRY could function as a standalone optogenetic module in mammalian cells. We therefore set out to test the feasibility of using DmCRY for optogenetic control in this context.

## Results

### Design of DmCRY-based light-inducible plasmids and confirmation of inducibility

In this study, we tested whether DmCRY can function as an optogenetic tool in mammalian cells. Previous work showed that DmCRY undergoes light-induced degradation in mammalian cells without the need for overexpression of any *Drosophila*-specific interactors (23). To harness this property, we fused DmCRY to the Tet repressor (TetR), a well-characterized regulator of gene expression (24). In its simplest configuration, this design enables light-dependent control of gene expression. The gene of interest is placed downstream of a regulatory element containing overlapping cytomegalovirus (CMV) promoter and tet operator (tetO) sites (TO). TetR dimers bind TO sequences to repress transcription, whereas tetracycline binding releases repression and restores transcription (25).

To generate this system, we inserted the TO sequence from pcDNA5/FRT/TO into the pcDNA4.Myc-HisA plasmid, enabling zeocin selection for stable cell line generation if necessary. We then expressed a DmCRY-TetR fusion protein alongside DmCRY-V5His to confirm its light-dependent degradation in mammalian cells (Fig. 1*A*). DmCRY-V5His was used as a representative of wild-type DmCRY, as done in many previous studies (26–28). Unlike prior work that used 366-nm illumination (23), we employed standard white light (or blue LED light, in some confirmatory experiments such as human portrait formation work), allowing implementation without specialized equipment. As expected, fusing TetR to DmCRY reduced degradation compared with wild-type representative DmCRY-V5His (*SI Appendix*, Fig. S1*A*). Attenuated degradation has also been reported in other DmCRY fusions such as DmCRY-APEX2 and DmCRY-TurboID (29). Our design goal was to achieve robust repression in the dark while minimizing leaky activation under ambient light. For better reproducibility, we constructed a single plasmid co-expressing DmCRY-TetR and the TO-controlled effector gene(s) (Fig. 1*B*). The expression of DmCRY-TetR transcripts was driven by the CMV promoter, and a WPRE (woodchuck hepatitis virus posttranscriptional regulatory element) was inserted to enhance transcript stability. This cassette (called “cc”) was flanked by MluI restriction enzyme recognition sites to allow insertion into different effector plasmids (except dCas9, see Methods).

**Figure 1.**
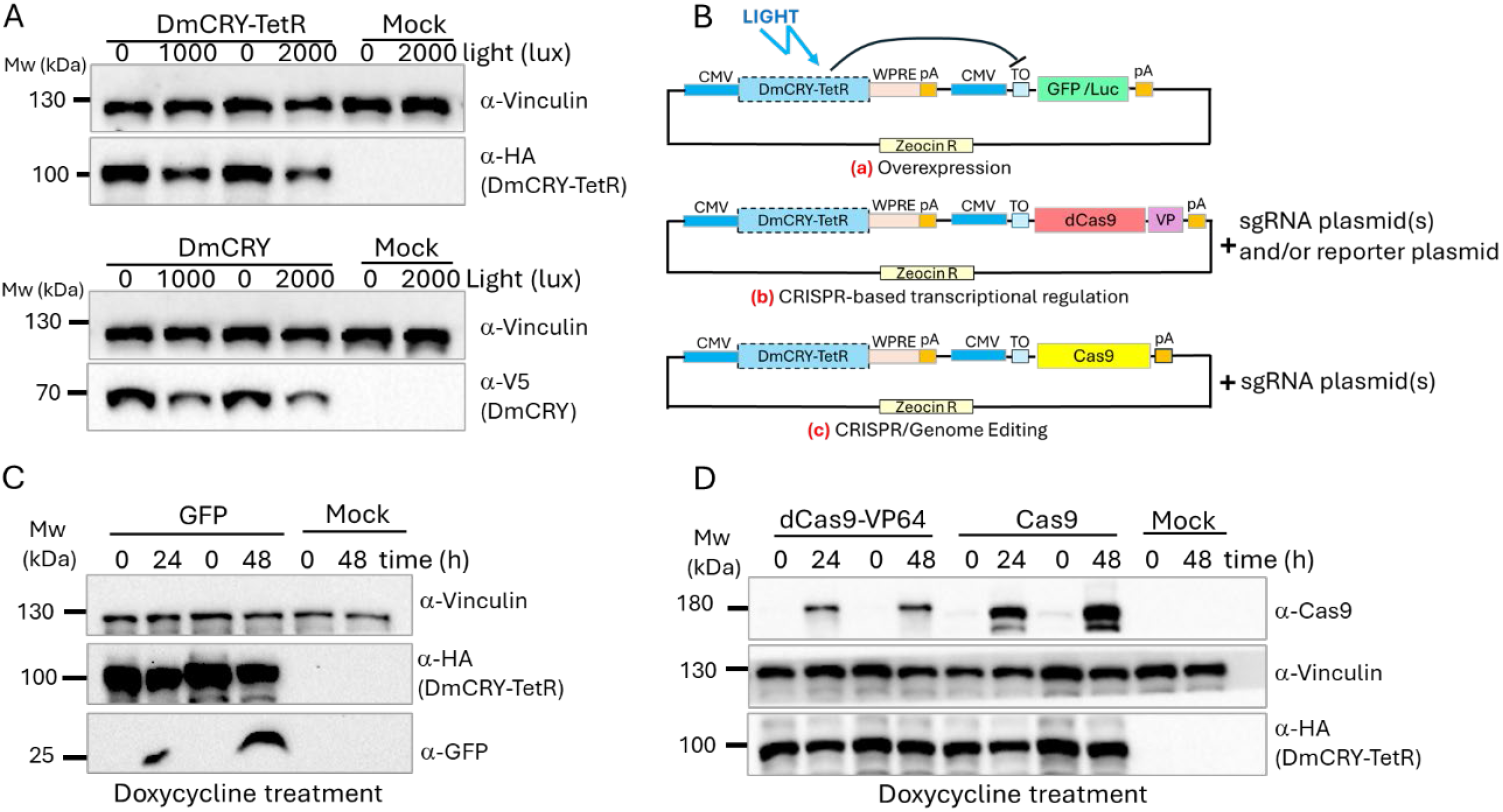
Light-dependent degradation of DmCRY in HEK293T cells and design of DmCRY-based inducible expression systems. (*A*) Immunoblots showing degradation of DmCRY-TetR and DmCRY-V5His (DmCRY) in HEK293T cells after exposure to 1,000 or 2,000 lux white light for 24 h. TetR fusion at the C terminus attenuated but did not abolish light-dependent degradation compared with DmCRY-V5His (wild-type reference). (*B*) Design of light- and tetracycline-inducible plasmids. Effector cassettes encoding GFP/Luc (a), dCas9 (b), or Cas9 (c) were placed downstream of TetR-responsive operator (TO) sequences. DmCRY-TetR, expressed from a separate cassette, binds TO sites to repress CMV-driven transcription. Removal of DmCRY-TetR by light-dependent degradation or tetracycline/doxycycline relieves promoter repression. VP, VP64 or VPR; Luc, luciferase cassette; pA, polyadenylation signal; R, resistance. (*C*) Doxycycline displaced DmCRY-TetR from OT sites, permitting CMV-dependent GFP expression and confirming TetR functionality in the DmCRY-TetR fusion. (*D*) Similarly, doxycycline displaced DmCRY-TetR from TO sites, permitting CMV-dependent expression of dCas9-VP64 and Cas9. Molecular weights (kDa) are shown to the left of blots. “Mock” indicates cells processed in parallel without plasmid transfection. Quantification and statistics for *A, C*, and *D* are given in *SI Appendix*, Figs. S1*A*–S1*C*, respectively.

Using this approach, we generated three systems: (a) Overexpression: a Green Fluorescent Protein (GFP) reporter to optimize light treatment conditions and image formation work, and a luciferase cassette (Luc) for image formation experiments. (b) CRISPR-based transcriptional regulation: a dCas9-based system co-expressed with sgRNAs and/or reporter plasmids. (c) CRISPR/genome editing: a Cas9 system for gene knockout when co-transfected with the relevant sgRNAs. To validate the functionality of TetR in DmCRY-TetR fusion, we tested whether it could repress target gene expression and retain tetracycline responsiveness. For these experiments, we used doxycycline, a more stable analogue of tetracycline, instead of tetracycline. Western blotting confirmed the induction of GFP expression upon doxycycline treatment (Fig. 1*C* and *SI Appendix*, Fig. S1*B*). Comparable expression patterns were also observed for dCas9 and Cas9 in the CRISPR-based transcriptional regulation and CRISPR/genome-editing systems, respectively (Fig. 1*D* and *SI Appendix*, Fig. S1*C*).

### Determination of optimal light dose, plasmid amount, and illumination time

After confirming that DmCRY-TetR effectively repressed CMV-OT activity and remained doxycycline-responsive, we next evaluated its responsiveness to light. Because prolonged light exposure at 366 nm can be cytotoxic to cells, we instead used visible light, which is less harmful and readily available in standard laboratories. White light exposure up to 2,000 lux was well tolerated by HEK293T cells, confirming its suitability for optogenetic applications (Fig. 2*A*). To identify optimal plasmid dose(s) for transfection, cells were transfected with increasing amounts of cc-pcDNA4-TO-GFP plasmid and illuminated with 2,000 lux white light for 24 h. GFP induction was clearly detectable with 250-500 ng transfected plasmid DNA with minimal background expression (Fig. 2*B*, and *SI Appendix*, Fig. S2*A*). Higher plasmid amounts could only increase fold-induction modestly, likely because both DmCRY-TetR and the target promoter resided on the same construct. We selected 500 ng as the working amount of transfected plasmid for subsequent experiments. We then tested varying light intensities (up to 3,000 lux which was the upper safe dose according to cytotoxicity results in Fig. 2*A*). GFP inductions increased in parallel to increased light doses (Fig. 2*C*, and *SI Appendix*, Fig. S2*B*). To balance the efficacy of target gene expression and cell viability, we adopted 1,000 lux as the standard illumination condition. Finally, we compared light-versus doxycycline-induced expression to estimate the maximal attainable induction. Whereas doxycycline fully relieved repression, light induced only partial activation (Fig. 2*D* and *SI Appendix*, Fig. S2*C*), consistent with incomplete DmCRY-TetR removal at subtoxic light doses. Importantly, this partial induction helps prevent inadvertent activation during cell handling, which can be particularly advantageous in genome-editing applications.

**Figure 2.**
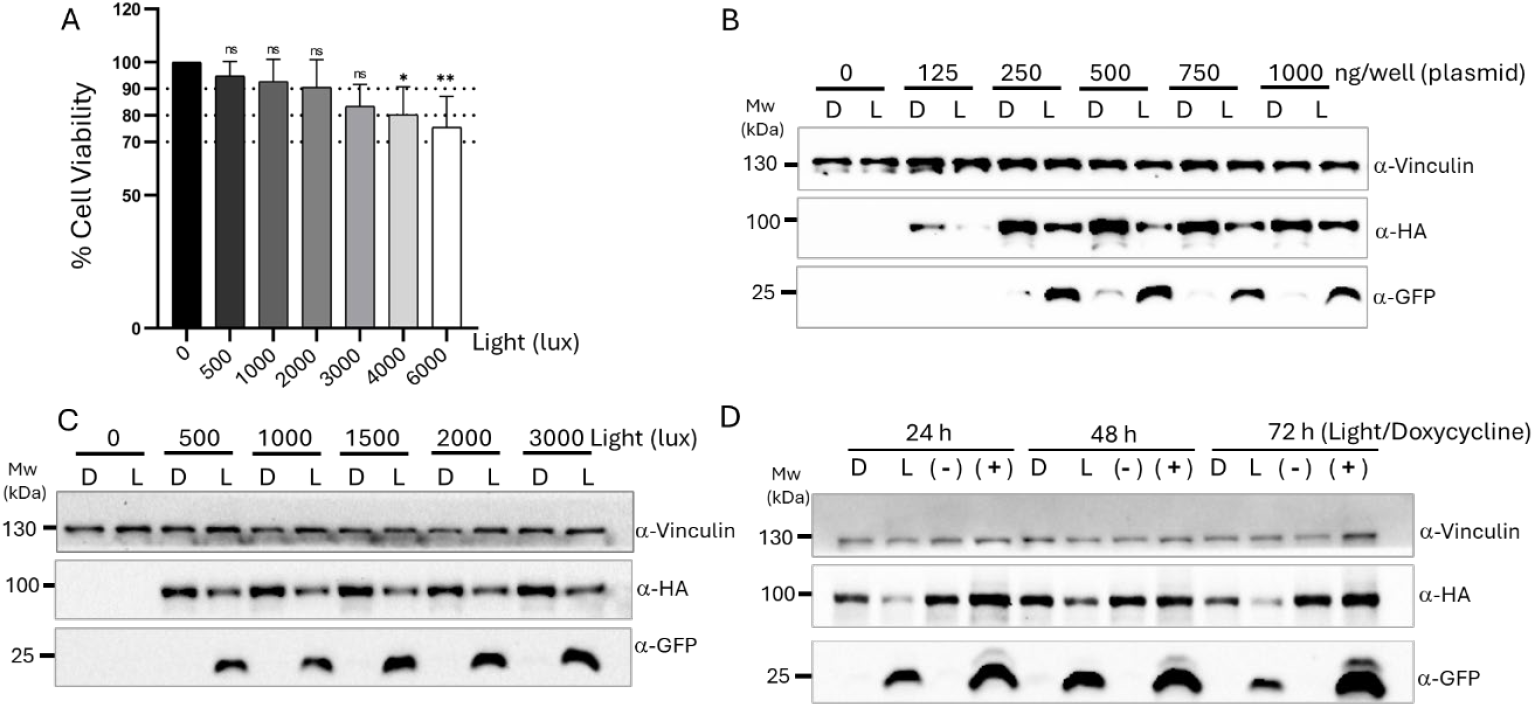
Optimization of light dose and transfected plasmid amounts. (*A*) The toxicity of white light was assessed at increasing intensities using an MTT assay. Cell viability after 48 h indicated tolerance to continuous exposure up to 3,000 lux. Statistical analysis was performed using one-way ANOVA with Tukey’s test in GraphPad Prism. Significance thresholds were defined as: ns < 0.1234, *p < 0.0332, **p < 0.0021, ***p < 0.0002, ****p < 0.0001. Results are presented as mean ± SD (n = 4). (*B*) GFP expression was tested at a subtoxic light dose (2,000 lux) with varying plasmid amounts. DmCRY-TetR levels were monitored by anti-HA immunoblotting. Optimal inducibility and expression were observed with 500 ng per well in a 24-well plate. (*C*) In the single-plasmid system, partial light-dependent degradation was sufficient to activate expression with minimal basal activity in the dark. Inducibility was therefore examined at subtoxic light doses. (*D*) Temporal control of effector expression by light versus doxycycline treatment. Induction kinetics were compared between doxycycline treatment and light exposure at different time points. D, dark; L, light (1,000 lux); (-), no doxycycline; (+), 1 μg/mL doxycycline. Quantitation and statistical analysis of blots were given in SI Appendix Figs. S2*A*-*C* for 2*B*, 2*C*, and 2*D*, respectively.

### Spatial control of light-induced expression and image formation by cells in dishes

Chemical inducers such as doxycycline provide robust activation but lack spatial precision. As a proof of concept, we tested whether our optogenetic system could achieve spatially restricted gene expression, a feature not possible with chemical regulators. To impose spatial light patterns, we generated photomasks by die-cutting polystyrene templates or by printing images on transparent acetate sheets, which were placed directly between cells and the light source during illumination. Transfected cells were illuminated through these masks, with the expectation that activated cells would form visible patterns through two reporter systems once light exposure ceased. Cells transfected with cc-pcDNA4-TO-GFP or cc-pcDNA4-TO-Luc and were exposed to light through photomasks of “L” letter or “CLK” letters which were made of polystyrene foams (*SI Appendix*, Fig. S3*A*). These letters were chosen to represent our laboratory, the Circadian Clock Lab. These photomasks were designed to cover 35-mm, 60-mm or 100-mm cell culture dishes. At the end of 24-hr white light exposure, fluorescence was recorded from the top of the plate after excitation with blue LED illumination, while luminescence was recorded after adding luciferin substrate, both recordings were captured with a CCD (Charge-Coupled Device) camera which was the same system used for immunoblotting detection (*SI Appendix*, Fig. S3*B*, top and lower panels, respectively). Images of simple patterns such as patterned circles, rectangular slits, or dots had been previously obtained with AtCRY-based optogenetic tools (6, 30), therefore we wanted to test if our system could offer more detailed image formation such as sketched images or even real human photos. For this purpose, we have printed images of a sketched clock, and a cat as well as a human portrait on the acetate sheets and up to six sheets were used as photomasks (*SI Appendix*, Fig. S3*C*). After the end of improvisation studies, cells transfected with cc-pcDNA4-TO-GFP were kept dark for 3 hours after a 24-h light exposure through photomasks. Because continuous light exposure causes bleaching of GFP signal, we allowed accumulation of newly synthesized GFP from CMV-TO-promoter from transcripts which were synthesized before illumination was stopped. Both GFP and luciferase images were presented in black and white (gray) and signal intensity-inverted formats, so it was easy to compare the image qualities. In Fig 3, top panel shows the GFP-images of a clock or a cat sketch and a human portrait (Figs. 3*A, 3B*, *and 3C, respectively*), while the lower panel shows luminescence images in the same settings (Figs. 3*D*, 3*E*, and 3*F*, respectively). The templates of the photomasks were given as insets on the top-right of each image. The imaging system could see only one 100-mm dish while taking fluorescence signal and could see only two 100-mm cell culture dishes at once. Fig. 3 shows cropped images of cell culture dishes taken at the same setting for each group. Therefore, at-once exposures of dishes were provided separately for a better comparison between light and dark samples as well as light-treated but not transfected mock cells (*SI Appendix*, Fig. S3*D*).

**Figure 3.**
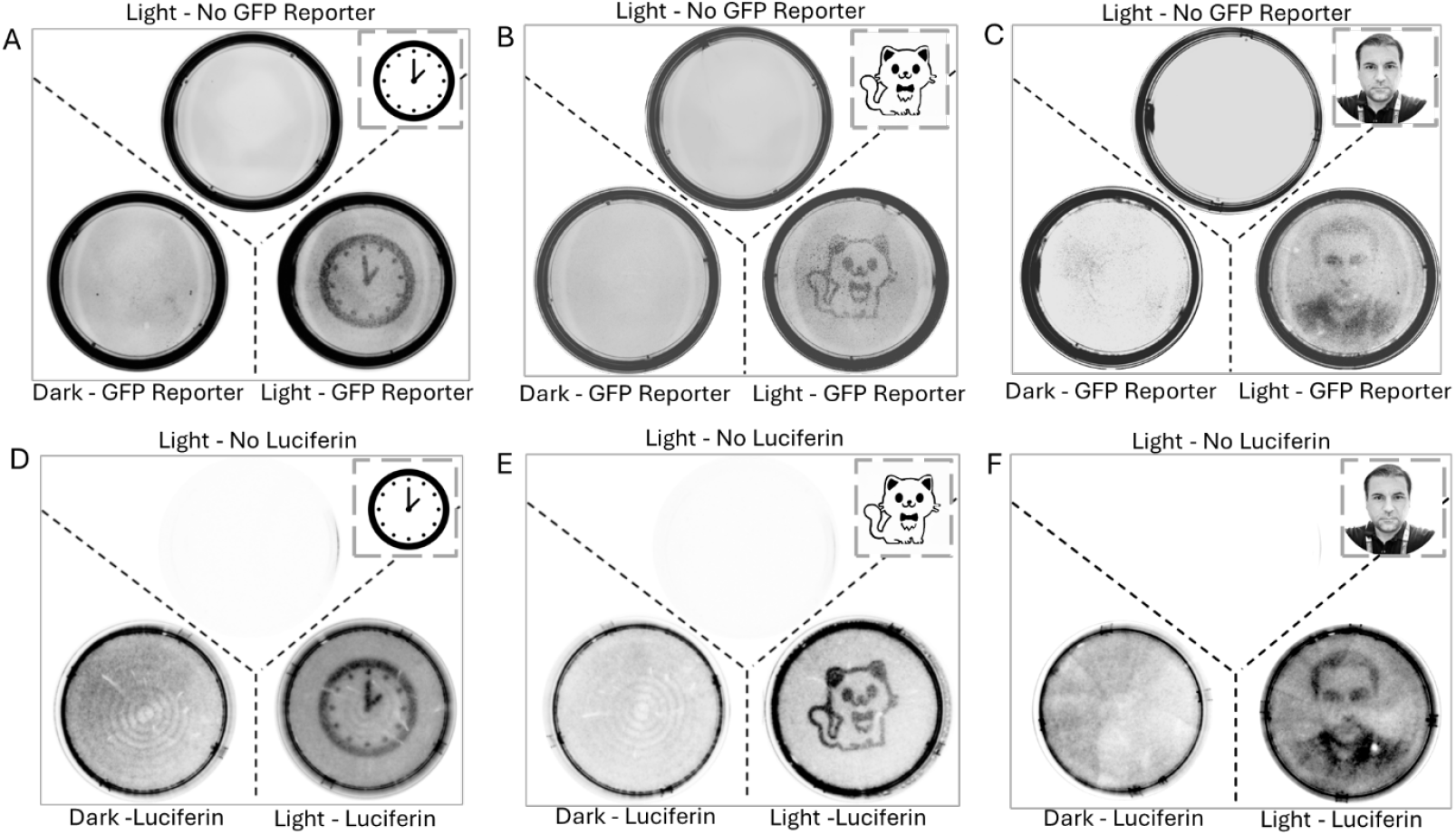
Spatial control of overexpression for image formation in cell culture by light. (*A*-*C*) Fluorescence imaging showing local control of overexpression in transfected cell populations. Cells were transfected with cc-pcDNA4.TO-GFP and exposed to light through the multiple layers of acetate paper sheets printed with a sketched clock (*A*), a sketched cat (*B*), or a human portrait (*C*). GFP signals were imaged across the entire dishes. Initial trials using polystyrene photomasks with the same reporter system are shown in *SI Appendix*, Fig. S3*A* and *B*. (*D*-*F*) Spatial control using a luciferase reporter under the same conditions. Cells were transfected with cc-pcDNA4.TO-Luc and exposed to light through multilayer acetate sheets printed with a sketched clock (*D*), a sketched cat (*E*), or a human portrait (*F*). Luciferase activity was imaged after the addition of luciferin. Templates of the sketched figures and portrait are shown as insets; all templates/photomasks are provided in *SI Appendix*, Fig. S3*C*. Because the imaging system could capture only one 100-mm dish at a time for fluorescence or two dishes at once for luminescence, the figure shows circular-cropped images taken under identical exposure settings. Partial views of all three dishes (for fluorescence) and whole two-dish combinations (for luminescence) are shown in *SI Appendix*, Fig. S3*D*.

As a further control to show images were specific but not artefact of light treatment, we transfected pcDNA4-TO-Luc plasmid for constitutive expression (which did not have “cc” to express DmCRY-TetR) and cc-pcDNA4-TO-Luc plasmid and exposed both to light side by side through “L” letter photomasks (*SI Appendix*, Fig. S3*E*). Comparison of images when both constitutively expressed and light-regulated luminescence had similar background signals revealed that only cc-pcDNA4-TO-Luc transfected cells formed a pattern while light exposure of constitutively expressed luciferase from pcDNA4-TO-Luc did not form a pattern which confirms that patterns formed were not due to a possible light effect on luciferase enzyme activity. Both fluorescence and luminescence approaches generated clear light-defined images even though luciferase system could generate superior images if we compare the clock images in Fig.3 *A* and *D*, demonstrating that DmCRY-TetR enables precise spatial control of gene expressions in mammalian cells. Larger images of single dish views or sketched human portraits, which were obtained during optimization with different photomasks or numbers of acetate sheets, were also presented as examples of light-induced spatial control of activity (*SI Appendix*, Fig. S3 *F* and *G*, respectively).

### Light-induced CRISPR-based transcriptional regulation

We next tested whether our system could be applied to transcriptional regulation of exogenous/endogenous genes. As an output module, we used dCas9-based transcriptional activation. In contrast to previous approaches that relied on AtCRY2-CIB1 interactions (30), our design expressed dCas9-VP64 under OT regulation. In this setup, sgRNAs directed dCas9-VP64 to a site upstream of a minimal promoter driving GFP expression. Light stimulation induced dCas9-VP64 expression from the cc-pcDNA4-TO-dCas9-VP64 plasmid, resulting in robust GFP activation from the reporter construct (Fig. 4*A*, and *SI Appendix*, Fig. S4 *A* and *B*). In our system, the synthesis of dCas9-VP64 protein after light exposure takes approximately one day, so there would be a delay from light exposure to reporter GFP induction by dCas9-VP64. Hence, we tested GFP reporter signal at 48 and 72 h following light treatment. To extend this strategy to endogenous gene control, we replaced VP64 with the more potent tripartite activator VP64-p65-Rta (VPR) (31). Transfection conditions were scale-down for 96-well plates to be able to use single-step RT-qPCR from cell lysates directly. Using sgRNAs targeting the CRY1 promoter, light stimulation successfully induced CRY1 expression via dCas9-VPR (Fig. 4*B*).

**Figure 4.**
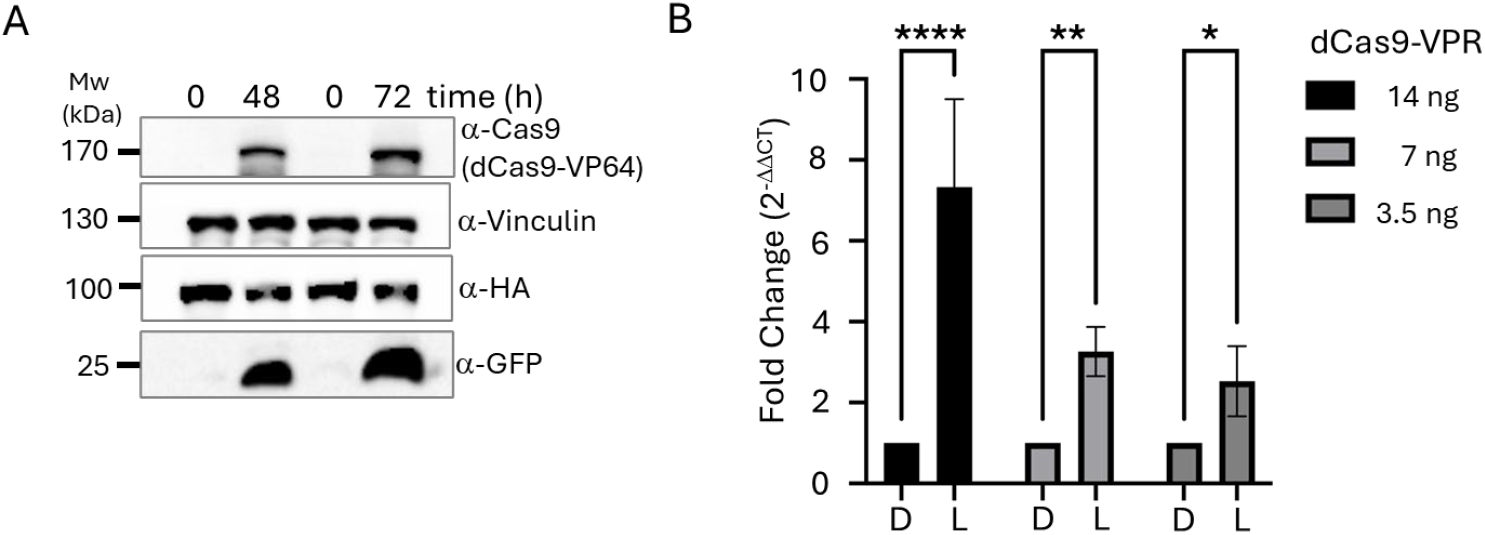
CRISPR-based regulation of gene expression by light. (*A*) Recombinant gene expression control through dCas9-VP64 was tested by monitoring GFP reporter expression. cc-pcDNA4.TO-dCas9-VP64 was co-transfected with the GFP reporter and the sgRNA plasmid targeting the reporter (see Methods). In this design, the expression of the effector (dCas9-VP64) occurs first and then the accumulated dCas9-VP64 subsequently activates GFP transcription. Therefore, only 48 h and 72 h samples were analyzed by Western blotting to assess reporter activation. Quantification and statistical analysis are shown in *SI Appendix*, Fig. S4. (*B*) Endogenous gene transcription control. Since VPR is reported to be more potent than VP64 for activating endogenous genes, HEK293T cells were co-transfected with cc-pCDNA4.TO-dCas9-VPR and lentiGuide-Puro-CRY1 plasmids. Induction was measured at the transcriptional level by qPCR. Statistical analyses were performed using two-way ANOVA followed by Tukey’s multiple comparisons test in GraphPad Prism. Significance thresholds were defined as: ns < 0.1234, *p < 0.0332, **p < 0.0021, ***p < 0.0002,****p < 0.0001. Results are presented as mean ± SD (n = 5).

### CRISPR/Cas9-mediated light-inducible genome editing

Finally, we tested whether our system could be extended to CRISPR/genome editing with CRISPR/Cas9 genome editing technique. Because knockout phenotypes require time to manifest, we followed an experimental design adapted from a previous study (32). As a control, constitutive Cas9 expression with a p53-targeting sgRNA plasmid efficiently reduced p53 protein levels, with nearly complete loss observed after one week (Fig. 5*A*). We then co-transfected cc-pcDNA4-TO-Cas9 with the same sgRNA plasmid (LentiGuide-Puro-p53) and monitored p53 expression after 5 and 6 days with or without light exposure. Light-treated cells showed a marked reduction in p53 protein compared with dark controls (Fig. 5*B*), and quantification confirmed a statistically significant decrease (Fig. 5*C*). These results demonstrate that the DmCRY-TetR system enables light-inducible genome editing in mammalian cells.

**Figure 5.**
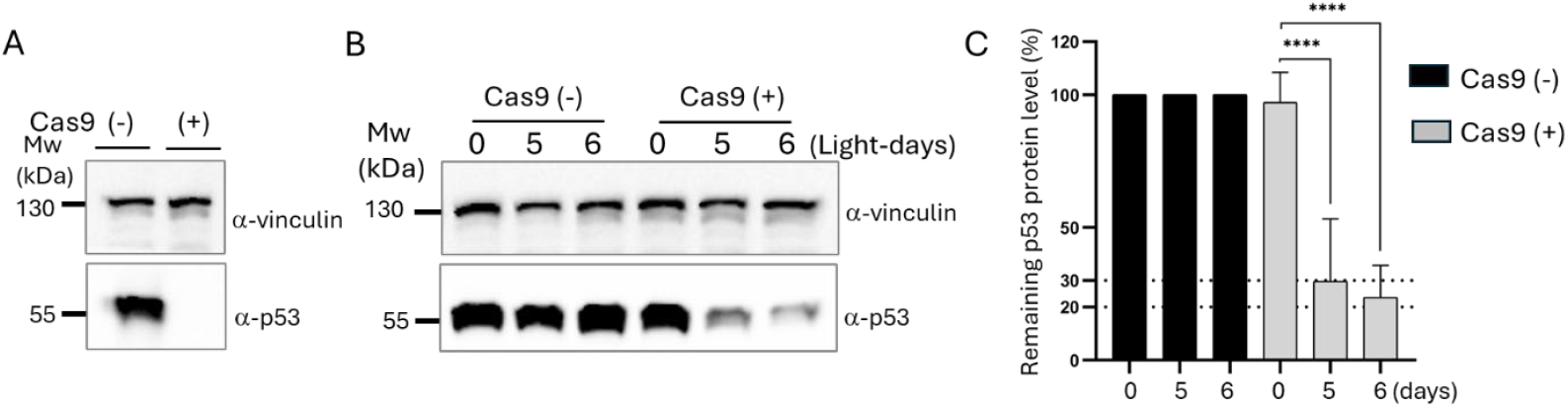
Light-controlled CRISPR/Cas9-mediated gene editing. (*A*) To validate the sgRNA can target *TP53* gene, HEK293T cells were co-transfected with pcDNA4.TO-Cas9 (constitutive Cas9 expression) and lentiGuide-Puro-p53 (expressing the sgRNA targeting *TP53*). Western blotting for p53 confirmed that Cas9 and the relevant sgRNA expression led to a detectable reduction in p53 protein levels. (*B*) Light-dependent Cas9 expression from cc-pcDNA4.TO-Cas9, together with sgRNA expression from lentiGuide-Puro-p53, enabled light-dependent knockout of *TP53*, as evidenced by reduced p53 protein levels in Western blotting. (*C*) Quantification of p53 reduction. p53 protein levels were compared between cells which were exposed to light 0, 5, and 6 days and normalized to vinculin. Statistical analyses were performed using one-way ANOVA followed by Tukey’s multiple comparisons test in GraphPad Prism. Significance thresholds were defined as: ns < 0.1234, *p < 0.0332, **p < 0.0021, ***p < 0.0002, ****p < 0.0001. Results are presented as mean ± SD (n = 5).

## Discussion

Although circadian photoreceptors from diverse taxa, including plants, insects, and vertebrate, contain stoichiometric flavin when expressed in heterologous hosts as an indication of their possible light-responsive functions (7, 9, 33), only AtCRY2 has been successfully adapted for optogenetic applications. This is largely due to its well-defined light-dependent interaction with CIB1 and its ability to oligomerize upon illumination. Even though DmCRY also has well-characterized light-dependent interactors, such as Jetlag and Timeless or recently identified ones (29), yet no optogenetic applications based on DmCRY have been reported except not fully successful trial of dimerization of an effector by fusing the same effector to DmCRY and Jetlag (34). We selected DmCRY over other alternatives such as type 4 CRYs (e.g., ZfCRY4), owing to our deeper mechanistic understanding of its properties, including its characteristic light-induced degradation in mammalian cells (23).

A central innovation of our work was to exploit this intrinsic degradation behavior of DmCRY by fusing it to TetR, enabling light-controlled repression without requiring additional *Drosophila* proteins. Importantly, TetR fusion attenuated the rapid degradation of DmCRY, improving system stability during cell handling. We further streamlined the system by constructing single plasmids encoding both the photosensory and effector modules, thereby facilitating modular customization for applications ranging from reporter induction to genome editing. System characterization demonstrated low basal expression, robust induction by white light intensities of more than 500 lux, and predictable saturation kinetics with modest plasmid amounts. Compared with doxycycline induction, light stimulation yielded lower absolute expression, reflecting the continuous replenishment of DmCRY-TetR from continuous expression even in the light. Nevertheless, the system’s capacity for spatial regulation represents a clear advantage over chemical inducers. Photomasking was previously used for simple pattern formations (30, 35). Using photomasks, we successfully generated precise patterns of gene expression, including recognizable human portrait images in cell culture dishes. We believe that this light signal intensity is transformed proportionally into reporter activation, as our design does not require multiple copies of the regulator regions for high expression. These findings highlight the utility of DmCRY-based systems for image formation and other applications requiring fine spatial control. Beyond reporter activation, we extended this system to transcriptional regulation using dCas9 effectors. While VP64 provided activation with the transfected reporter with 8 copies of sgRNA binding sequences, substitution with the tripartite activator VPR enabled robust light-inducible activation of the endogenous CRY1 gene. Finally, we confirmed genome editing capability by coupling Cas9 expression to light control, achieving efficient reduction of p53 protein due to knockout in HEK293T cells without antibiotic selection. Together, these results illustrate the versatility of DmCRY-TetR in regulating a broad range of outputs. A key advantage of DmCRY is its functional independence from native *Drosophila* interaction partners, which simplifies deployment in mammalian systems. Nonetheless, additional opportunities remain. For example, modifications can be introduced to the DmCRY C-terminus or the conserved tryptophan triad (36) to dissect the activation mechanism of the tool and to lower the background expression. Such variants could expand the repertoire of DmCRY-based optogenetic applications, enabling the rational tuning of degradation dynamics or interaction specificity.

In summary, we introduce DmCRY as a novel optogenetic module enabling temporal and spatial control of gene expression, transcriptional regulation, and genome editing. We envision that red, blue, and green (RGB) reporters, activated by corresponding photoreceptors, could one day be combined to generate live, full-color human portraits in cell culture. This platform broadens the scope of clock-based optogenetics beyond AtCRY2 and lays the foundation for future DmCRY-based tools with enhanced precision and versatility.

## Materials and Methods

### Plasmids

For this study, we generated three classes of plasmid systems. 1-For expression of reporters such as Green Fluorescent Protein (GFP) and luciferase (Luc), 2-For transcription regulation through nuclease dead Cas9 (dCas9), and 3-For gene editing through Cas9. The reporters (GFP and Luc) or effectors (dCas9 and Cas9) were controlled by a sensor system within modular vectors assembled for specific purposes. A tetracycline repressor sequence (TetR) fused to a nuclear localization signal (NLS) coding sequence (TetR-NLS) was amplified from pLVUT-tTR-KRAB plasmid (Addgene no. 11651) (37) using oligonucleotides with 5’ TTGTCTAGAATGGCTAGATTAGATAAAAG 3’ and 5’ TTACCCGGGGGCACCACCGCCGTCGACCTT 3’ sequences. Then PCR product was digested with XbaI and SalI restriction enzymes and cloned into the XbaI-XhoI site of pLenti III-HA vector (Applied Biological Materials Inc., Richmond, Canada) to generate pLenti-III-TetR-NLS-HA plasmid. The Drosophila Cryptochrome (DmCRY) coding sequence from pAc5.1.DmCRY-V5His plasmid (38) was then inserted into the EcoRI-XbaI sites of pLenti-III-TetR-NLS-HA, yielding pLenti-III-DmCRY-TetR-NLS-HA. The DmCRY-TetR-NLS-HA cassette was excised by NheI-PmeI digestion and transferred into the NheI-EcoRI sites of pCMV-3Tag6 (Agilent Technologies, cat no. 240200) to generate pCMV-3Tag6-DmCRY-TetR-NLS-HA. A SalI-WPRE-SalI fragment from LentiCRISPR v2 plasmid (Addgene no. 52961) (39) was excised and ligated into the SalI site of pCMV-3Tag6-DmCRY-TetR-NLS-HA; correctly oriented clones were identified by XbaI-SacII restriction enzyme analysis. The CMV-DmCRY-TetR-NLS-HA-WPRE sequence was subsequently amplified from this construct using oligonucleotides with 5’ GTAACGCGTCATAGCCCATATATGGAGTTC 3’ and 5’ CAATTTACGCGTTCCCCAGCAT 3’ sequences. The amplified sequence was named as “cc*”* module and digested with MluI restriction enzyme and inserted into MluI sites of the effector plasmids, which had TetR responsive elements.

The plasmids for the effector expression were also constructed in a modular order. MluI-HindIII fragment of pcDNA5.FRT/TO (Thermo Fisher Scientific, Waltham, Massachusetts, USA, cat no: V652020) was transferred into with MluI-HindIII of pcDNA4/myc-His A, (Thermo fisher scientific, cat no: V863-20) to obtain pcDNA4.TO/MycHisA. This plasmid has tetR responsive sequences downstream of the CMV promoter. To obtain pcDNA4.TO-EGFP, HindIII-NotI fragment from pEGFP_N2 (Clontech, Mountain View, CA, USA) was transferred into HindIII-NotI site of pcDNA4.TO/MycHisA. To obtain pcDNA4.TO-Cas9, Cas9 was transferred from pCW-Cas9 plasmid (Addgene no. 50661) (40). For this pcDNA4.TO/MycHisA was digested with HindIII, overhangs were filled with Q5 High-Fidelity DNA Polymerase (New England Biolabs, Ipswich, MA, USA, cat no: M0491S) and then digested with BamHI restriction enzyme. Meanwhile, pCW-Cas9 was digested with NheI and filled with Q5 polymerase and then digested with BamHI restriction enzyme. Directional cloning of Cas9 into the Blunt and BamHI site of pcDNA4.TO/MycHisA resulted in pcDNA4.TO-Cas9 plasmid. pcDNA4.TO-dCas9-VP64 was obtained through a multi-step cloning strategy. dCas9 was amplified from pcDNA3.1-CibN-dCas9-CibN (Addgene no. 60553) (30) using oligos 5’ GATCTCGAGCATCACGATTACAAGGATG 3’ and 5’ CCCTCTAGAAACTTTGCGTTTCTTTT 3’ sequences and inserted into XhoI-XbaI site of pcDNA4.MycHisA to obtain pcDNA4.dCas9. A linker was generated using annealed oligos with 5’ GGCCGCAGATCTCGACACCATGGTGC 3’ and 5’ TCGAGCACCATGGTGTCGAGATCTGC 3’ sequences and inserted into the NotI-XhoI site of pcDNA4.dCas9 to generate and ATG with frame with the dCas9 coding sequence. Then NheI-VP64-PmeI fragment from pcDNA3.1-Cry2FL-VP64 plasmid (Addgene no. 60554) (30) was inserted into the XbaI-PmeI site of this plasmid to obtain pcDNA4-dCas9-VP64. From this plasmid, BglII-dCas9-Vp64-XbaI was transferred to BamHI-XbaI site of pcDNA4.TO/MycHisA to obtain pcDNA4.TO-dCas9-Vp64. cc-module (CMV-DmCRY-TetR-NLS-HA-WPRE) was inserted into MluI sites of pcDNA4.TO/GFP and pcDNA4.TO-Cas9 to obtain cc-pcDNA4.TO-GFP and cc-pcDNA4.TO-Cas9, respectively. Right oriented ones compared to CMV.TO-GFP or CMV.TO-Cas9 cassettes were selected by restriction enzyme analysis. Because dCas9 had MluI site, we could not use MluI cloning to obtain cc-pcDNA4.TO-dCas9-VP64. Instead, we transferred a 4721 bp-BglII fragment from cc-pcDNA4.TO-Cas9 into BglII site of pcDNA4.TO-dCas9-VP64 and selected the right oriented one clone by restriction enzyme analysis to obtain cc-pcDNA4.TO-dCas9-VP64. The gRNA-eGFP-Reporter (Addgene no. 60719) which expresses guide RNA used to activate its reporter plasmid pGL3-Basic-8x-gRNA-eGFP (Addgene no. 60718) were obtained from Addgene and was part of LACE system (30). pcDNA4.TO-dLuc was generated by a two-step cloning process. The NcoI-XbaI fragment encoding Luc protein from pGL3.Basic was transferred into NcoI-XbaI site of pGL3-Basic with NcoI and XbaI site of pGL3-Basic-8x-gRNA-eGFP to obtain pGL3-Basic-8x-Luc. Then HindIII-XbaI fragment from this plasmid was transferred into HindIII-XbaI site of pcDNA4.TO/MycHisA to obtain pcDNA4.TO-Luc. Then cc module was inserted into MluI site of pcDNA4.TO-Luc to obtain cc-pcDNA4.TO-Luc. To target *TP53* gene via with Cas9, sgRNA was expressed from the lentiGuide-Puro plasmid (Addgene no. 52963) (39). To obtain *TP53* sgRNA expressing plasmid, p53 Sense 5’ CACCGCCATTGTTCAATATCGTCCG 3’ and p53 Antisense 5’ AAACCCGGACGATATTGAACAATGGC 3’ oligos were annealed and inserted into BsmBI site of the lentiGuide-Puro plasmid. To activate endogenous *CRY1* transcription using dCas9, three different CRY1 sgRNAs targeting CRY1 promotor region were generated from three sets of annealed oligos with the sequences as follow: CRY1_P1 Sense 5’

CACCGAGAGCGCACGCGCGCCTAA 3’ and CRY1_P1 Antisense 5’

AAACTTAGGCGCGCGTGCGCTCTC 3’; CRY1_P2 Sense 5’

CACCGGTTTGGACTTGAAATGTACG 3’ and CRY1_P2 Antisense: 5’

AAACCGTACATTTCAAGTCCAAACC 3’; CRY1_P3 Sense 5’

CACCGCAGGATTGAAGCACATGGTT 3’ and CRY1_P3 Antisense: 5’

AAACAACCATGTGCTTCAATCCTGC 3’. Then annealed oligos were inserted into BsmBI-digested lentiGuide-Puro plasmid. Unless otherwise stated all enzymes used for cloning were purchased from Thermo Fisher Scientific.

### Cell Lines

HEK293T cells were obtained from the American Type Culture Collection (ATCC, Rockville, MD) and maintained in a humidified incubator at 37 °C with 5% CO_2_. Cells were cultured in Dulbecco’s Modified Eagle Medium (DMEM; Thermo Fisher Scientific, cat no. 41966029) supplemented with 1% nonessential amino acids (Thermo Fisher Scientific, cat no. 11140035), 100 U/mL penicillin and 100 µg/mL streptomycin (Thermo Fisher Scientific, cat no. 15140122), and 10% fetal bovine serum (FBS; Thermo Fisher Scientific, cat no. A5256701). For light treatment experiments, culture media were switched to CO2-independent medium (Thermo Fisher Scientific, cat no. 18045088) supplemented with 1% nonessential amino acids, 100 U/mL penicillin and 100 µg/mL streptomycin, 200 mM L-glutamine (Thermo Fisher Scientific, cat no. 25030024) and 10% FBS. Following medium exchange, cells were incubated under atmospheric CO_2_ conditions in a temperature-controlled incubator (FOC 120E Connect Cooled Incubator, Velp Scientifica, NY, USA). To minimize potential heating from local light exposure, during light exposure experiments cells were incubated at 35 °C to ensure that the effective sample temperature did not exceed 37 °C, which could otherwise cause stress or toxicity. Dark controls were also exposed to the light source but shielded with aluminum foil, ensuring that both light- and dark-treated samples experienced the same local thermal conditions.

### Light Treatment Experiments

Cells were seeded at 70–80% confluency in cell culture plates or dishes and transfected with the respective plasmids after overnight incubation in complete DMEM. Transfections were performed according to previously optimized protocols with small modifications (41, 42). For 24-well plates, up to 1 μg plasmid DNA was dissolved in 40 μL of complexation buffer (100 mM NaCl, 10 mM Tris·HCl, pH 7.5, 0.5 mM EDTA). Polyethylenimine (PEI; 1 mg/mL; Sigma, cat no. 408727) was added at a 3:1 weight ratio (PEI:DNA). The mixture was pipetted gently three times, incubated for 15 min at room temperature, combined with 1.5 mL of complete medium, and distributed into two wells (700 μL per well, in duplicate plates).

For 96-well plates (*see Fig. 4B*), volumes were scaled down 6-fold. Each well received ∼ 160 ng plasmid DNA (50 ng for each of 3 different lentiGuide-Puro-CRY1 plasmids and up to 14 ng of cc-pcDNA4.TO-dCas9-VPR plasmid) and dissolved in 7 mL of complexation buffer, mixed with 0.5 mL PEI. After incubation, the transfection complexes were diluted with 200 mL complete medium, and 100 mL was added per well (in duplicate for light and dark treatments). For photomasking experiments in 35-mm, 60-mm and 100-mm culture dishes, 3, 6 or 9 μg plasmid DNA was mixed with 200, 400 or 750 μL of complexation buffer, respectively, and diluted in 3, 7 or 18 mL of complete DMEM and distributed equally in two plates. In all cases, the 3:1 PEI:DNA ratio was maintained.

After 24 h, the medium was replaced with the CO2-independent medium, and plates were randomly assigned to light or dark treatment groups. For light exposure, cells were placed in an incubator without CO_2_ supplementation and illuminated with a dimmable LED desk lamp (Touch Eye Protection, TT-DL13 model, TaoTronics) or Light Therapy Lamp (UV-Free Sun Lamp with 660nm Red Light and 12000 Lux Bright White Light, GQP, UPC: 683404119938, China) using white light option at given light levels (in lux units) unless specified otherwise. Blue light was given using a Sunset Lamp Projector (Microcase, model no. 6805). Dark controls were covered with aluminum foil. All experiments were performed using white light; however, some experiments were repeated with blue light source (450 nm) at a rate of 2 milliwatt·cm^-2^ especially for a few formation experiments which is stated in the figure legends. A glass plate was positioned between the samples and the light source. When different light irradiance was required, it was achieved either by adjusting the distance from the light source or by placing neutral density filters between the samples and the light source. Illuminance at the plate plane was measured with Fisherbrand™ Traceable™ Dual Display Light Meter (Fisher Scientific, cat. no. 06-662-64) and verified using AquaHorti AH-300 light meter (AquaHorti). Illuminance is reported in lux for readability. Blue-band irradiance (400–500 nm) was measured with a Solarmeter Visible Blue Light Meter Model 9.4 (Solarmeter cat no. 9.4). Measurements were taken at several positions across the plate. Spectral properties and light measurement details are given in *SI Appendix*, Fig. S5.

### Cell viability (MTT) assay

HEK293T cells were seeded at 7.5×10^3^ cells per well (of a 96-well plate) in complete CO2-independent medium (with a final volume 100 μL/well) and allowed to adhere for 24 h at 37 °C. Cells were then exposed to defined light doses by varying exposure time at a fixed, measured irradiance at the plate plane. Dark controls were included for each light treatment. After light treatment, MTT (thiazolyl blue tetrazolium bromide; stock 5 mg/ mL in 1X PBS) was added to each well to a final concentration of 0.5 mg/mL (e.g., add 10 μL MTT stock to 90 μL medium per well) and plates were incubated for 3 h at 37 °C, protected from light. The supernatant was carefully removed, 100 μL DMSO was added to dissolve formazan crystals, and plates were shaken at room temperature for 15 min. The optical density was measured at 570 nm with a Varioskan Flash multi-plate reader (Thermo Scientific, LS-5250040). Background (cell-free) blanks and vehicle controls were subtracted. Each experiment was repeated four times independently.

### Western blotting

Western blotting was performed with small modifications from a previously published protocol (41). Briefly, cells were harvested and lysed in radioimmunoprecipitation assay (RIPA) buffer (50 mM Tris-HCl pH 7.5, 150 mM NaCl, 1% IGEPAL and 0.15% Sodium Dodecyl Sulfate) supplemented with a protease inhibitor cocktail (Merck KGaA, cat. no. S8830-2TAB). Equal amounts of protein (50 μg) in cell lysates were separated by SDS-PAGE, and transferred to nitrocellulose membranes (Bio-Rad, cat no. 162-0115). Membranes were blocked in the blocking solution of 1X TBS-T (150 mM NaCl, 20 mM Tris-HCl, pH 7.6, 0.08% Tween-20) containing 5% nonfat dry milk and incubated overnight at 4 °C with primary antibodies diluted in the blocking solution. The following antibodies were used: anti-HA (Cell Signaling Technology, 3724S; 1:3,000), anti-V5 (Thermo Fisher Scientific, R96025; 1:5,000), anti-p53 (Cell Signaling Technology, 2524; 1:2,000), anti-Cas9 (Merck, SAB4200701, 1:3000) and anti-vinculin (Thermo Fisher Scientific, 14-9777-82; 1:2,000). Dilution factors are indicated after the catalog numbers.

After four washes in 1X TBS-T, membranes were incubated for 1 h at room temperature with horseradish peroxidase (HRP)-conjugated secondary antibodies (Advansta Inc, CA, USA): anti-mouse HRP (cat no. R-05071-500; 1:10,000) or anti-rabbit HRP (cat no. R-05072-500; 1:10,000). Following four 1X TBS-T washes, chemiluminescence was detected with Glossy Femto HRP-Substrate (Nepenthe, Kocaeli, Türkiye, cat no. B0512043C1) and images were captured using ChemiDoc XRS+ system (Bio-Rad). Molecular weight standards were determined using PageRuler protein ladder (Thermo Fisher Scientific, cat. no. 26616).

### Cellular Imaging

Cells were transfected with cc-pcDNA4.TO-GFP or cc-pcDNA4.TO-Luc in 35-mm, 60-mm or 100-mm dishes as described above and incubated overnight. For light treatment, the medium was replaced with CO2-independent medium, and cells were exposed for 24 h to white light through 2-cm thick polystyrene foam (black-painted) templates with letters carved into them or acetate sheets with figures or photos printed on. At the end of light treatment, images from cell plates were acquired using appropriate image analyzer equipment. Fluorescence (GFP) signals were recorded on an Azure Biosystems imager (Azure Biosystems, c600, Dublin, CA, USA) using a green filter (Kodak, Green 58 Wratten Color Filter, cat no:53-701). For bioluminescence imaging, 150 µg/mL luciferin substrate (Promega, Madison, WI, USA; cat. no. E1601) was added to the culture medium, and luminescence was captured using an Azure Biosystems imager.

### Real-time quantitative PCR analysis

Cells were seeded in 96-well plates (in duplicate) and exposed to light for 24 h, with aluminum foil–covered wells serving as dark controls. Transcript levels were analyzed using the Luna Cell-Ready One-Step RT-qPCR Kit (New England Biolabs, Ipswich, MA, USA, cat no. E3030S). qPCR results were analyzed following a previous report (43). *CRY1* expression levels were normalized to those of *GAPDH*. The oligonucleotide sequences were as follows: *CRY1* forward, 5′ ACGAGGGGACCTGTGGATTA 3′; reverse, 5′AGGACAGGCAAATAACGCCT 3′; and *GAPDH* forward, 5′ TGCACCACCAACTGCTTAGC 3′; reverse, 5′ACAGTCTTCTGGGTGGCAGTG3′.

### Statistical Analysis

Statistical analyses were performed using GraphPad Prism 7 (GraphPad Software, La Jolla, CA, USA). Quantitative data are reported as the mean ± SD from at least three or more independent experiments. The specific statistical tests and sample sizes (n) are provided in the relevant figure legends.

## Ethics Statement

This study did not involve human participants or animals. The human images presented in Fig. 3, and *SI Appendix*, Fig. S3 is a photograph of one of the authors, who provided written informed consent for its use. No additional human subjects were involved.

## Use of AI Software

ChatGPT was used to improve the wording of some paragraphs, but not to generate new content.

## Acknowledgments

This study was financially supported by a research grant from the Health Institutes of Türkiye (TÜSEB) with grant number 2022-B-03-24282 to Nuri Ozturk. We thank Central Research Laboratory Application and Research Center (GTU-MAR), Gebze Technical University, Gebze, 41400, Kocaeli, Türkiye for allowing us to use their infrastructure facilities. We are also grateful to our undergraduate project students, Irem Unal and Inci Yerli, for their assistance with general laboratory work during this study, and to our former lab member, M. Serdar Koca (Universidad de Granada), for his guidance in statistical analysis.

## Notes

### Competing Interest Statement

The authors have declared no competing interest.

