## Supplementary Information (SI) for "Single-component optogenetic control enables human portrait formation in cells"

Figure S1

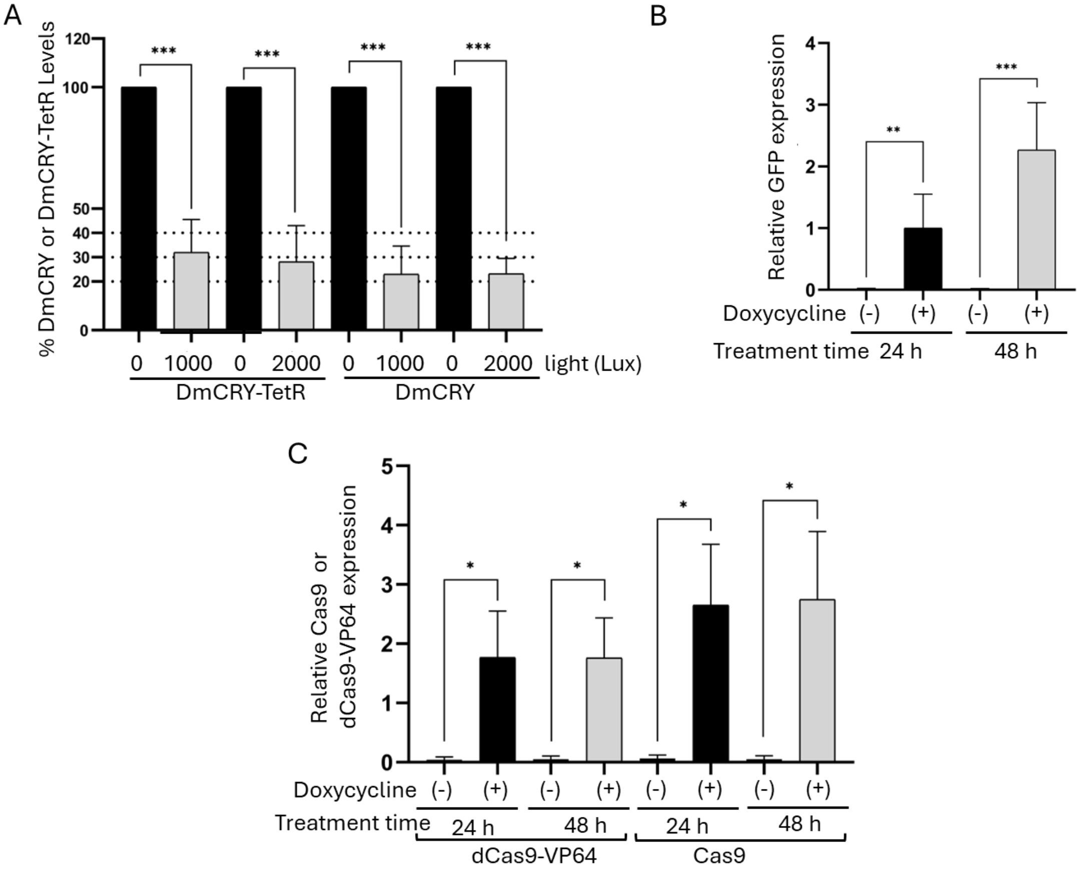


**Fig. S1.** Quantification and statistical analysis of DmCRY degradation and effector induction following doxycycline treatment. (*A*) Light-dependent degradation of DmCRY-TetR and DmCRY-V5His (DmCRY) in HEK293T cells was detected using anti-HA and anti-V5 antibodies, respectively, and normalized to vinculin levels. Band intensities from Western blot images (n = 6) were quantified and presented as mean ± SD. Differences between dark (0 lux) and illuminated samples (1,000 and 2,000 lux) were analyzed using a two-tailed unpaired Student's t-test in Graphpad Prism. (*B*) GFP induction (n = 5) and (*C*) dCas9-VP64 and dCas9 induction (n = 3) upon doxycycline treatment were quantified from Western blot images and presented as mean ± SD. Statistical significance between solvent (-) and doxycycline (+) samples was determined using a two-tailed unpaired Student's t-test in GraphPad Prism. Statistical significance was defined as: ns<0.12, *p<0.033, **p<0.002, ***p<0.0001. Images given in the main manuscript files, Figs 1*A*, 1*C*, and 1*D* are the representative ones which are used for quantitation in Figs S1*A*, S1*B*, and S1*C*, respectively.

Figure S2

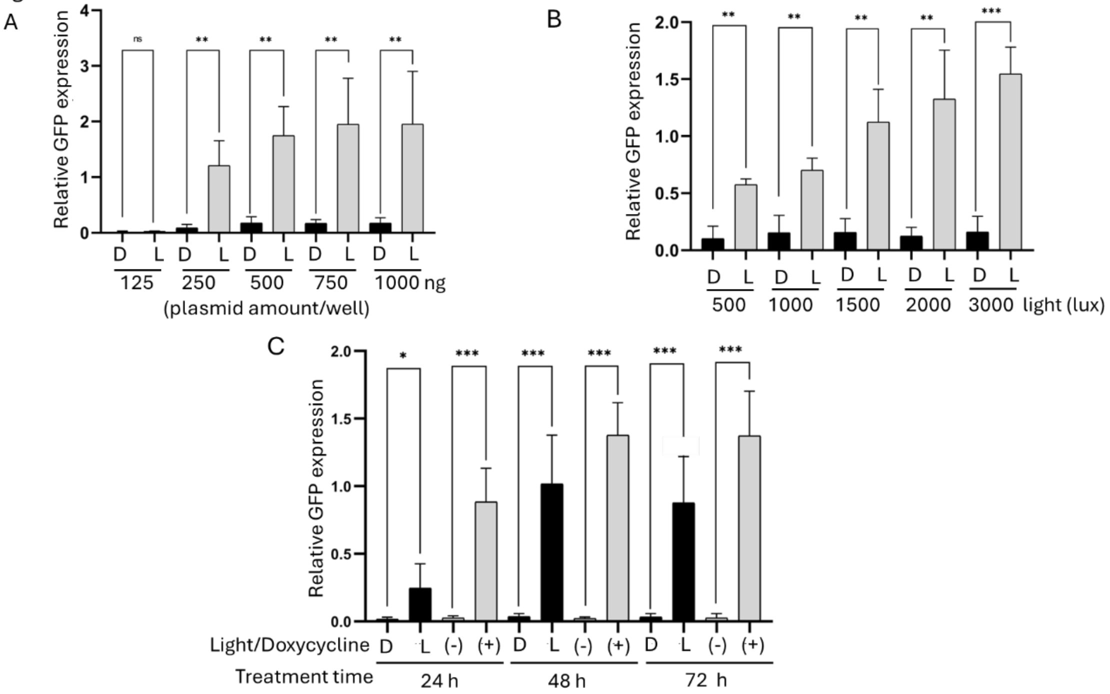


**Fig. S2.** Quantification and statistical analysis of GFP expression following light or doxycycline treatments. (*A*) GFP expression was assessed at a subtoxic light intensity (2,000 lux) with varying plasmid amounts to evaluate expression strength and inducibility. Band intensities from immunoblots (n = 4) were quantified and presented as mean ± SD. Differences between dark (0 lux) and illuminated (2,000 lux) samples were analyzed using a two-tailed unpaired Student's t-test in GraphPad Prism. (*B*) Light dose–dependent induction of GFP expression. Band intensities from Western blots (n = 3) were quantified and presented as mean ± SD. Statistical differences between each dark (0 lux) and light-treated samples (500–3,000 lux) were determined using a two-tailed unpaired Student's t-test in GraphPad Prism (*C*) Temporal control of effector expression by light versus doxycycline treatment. GFP induction kinetics were compared between light exposure and doxycycline treatment at different time points. Band intensities from Western blots (n = 5) were quantified and presented as mean ± SD. Statistical differences between dark (D) and light (L) or between solvent (-) and doxycycline (+) samples were determined using a two-tailed unpaired Student's t-test in Graphpad Prism. The levels of statistical significance were defined as: ns<0.12, *p<0.033, **p<0.002, ***p<0.0001. Images given in the main manuscript files, Figs 2*B*, 2*C*, and 2*D* are the representative ones which are used for quantitation in Figs. S2*A*, S2*B*, and S2*C*, respectively.

Figure S3

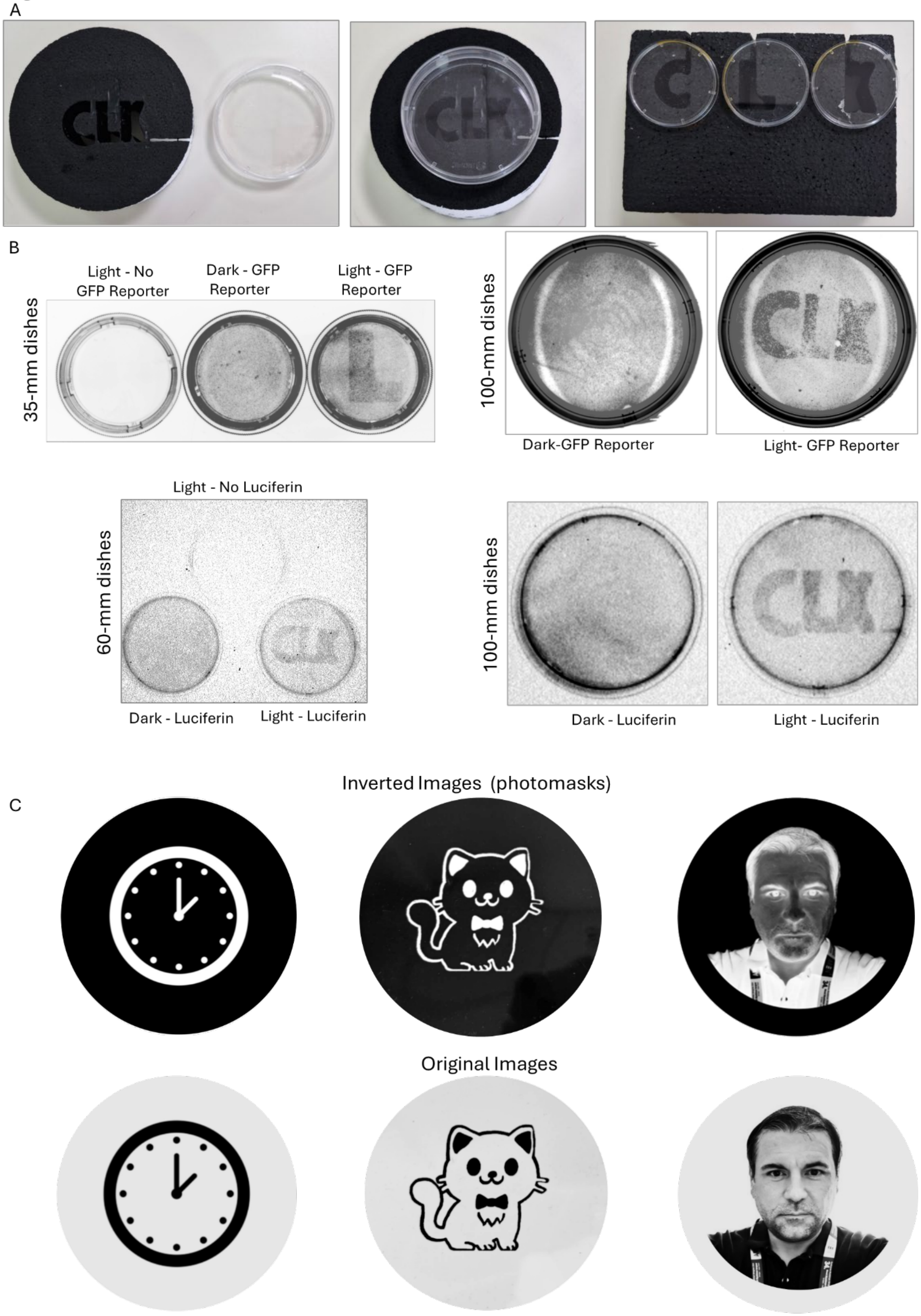
A
B
Light - No GFP Reporter
Dark - GFP Reporter
Light - GFP Reporter
35-mm dishes
100-mm dishes
Dark-GFP Reporter
Light- GFP Reporter
Light - No Luciferin
60-mm dishes
Dark - Luciferin
Light - Luciferin
100-mm dishes
Dark - Luciferin
Light - Luciferin
C
Inverted Images (photomasks)
Original Images

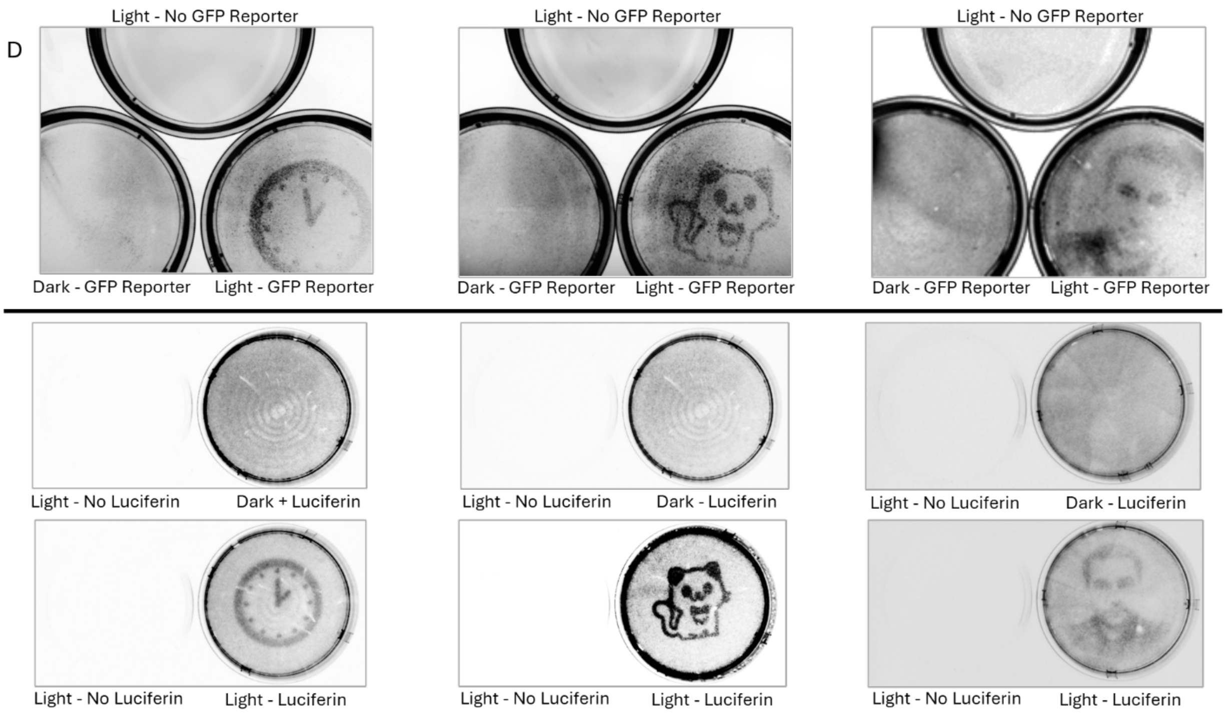
D
Light - No GFP Reporter
Dark - GFP Reporter
Light - GFP Reporter
Light - No GFP Reporter
Dark - GFP Reporter
Light - GFP Reporter
Light - No GFP Reporter
Dark - GFP Reporter
Light - GFP Reporter
Light - No Luciferin
Dark + Luciferin
Light - No Luciferin
Dark - Luciferin
Light - No Luciferin
Dark - Luciferin
Light - No Luciferin
Light - Luciferin
Light - No Luciferin
Light - Luciferin
Light - No Luciferin
Light - Luciferin

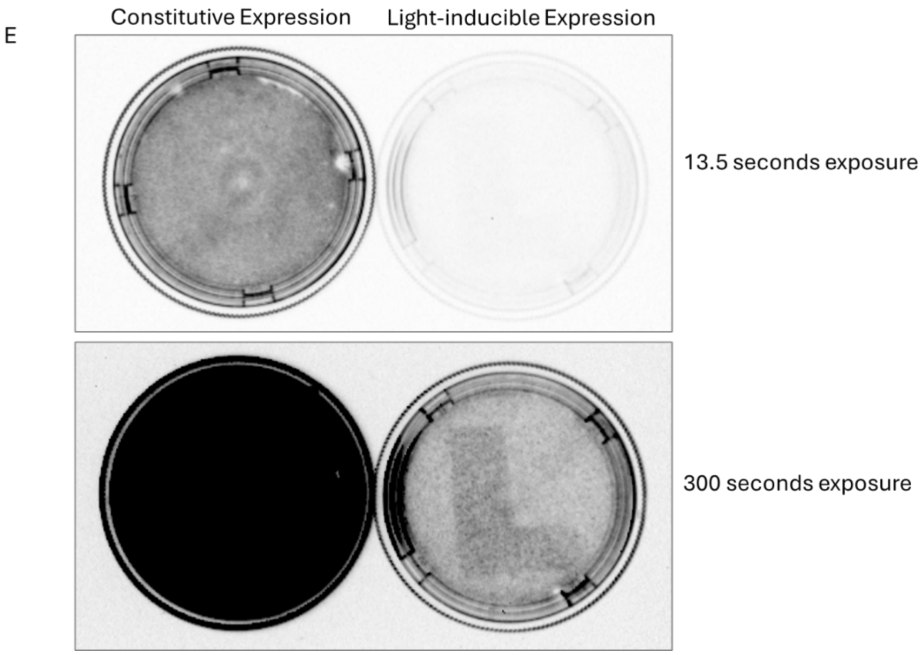
E
Constitutive Expression
Light-inducible Expression
13.5 seconds exposure
300 seconds exposure

F

G

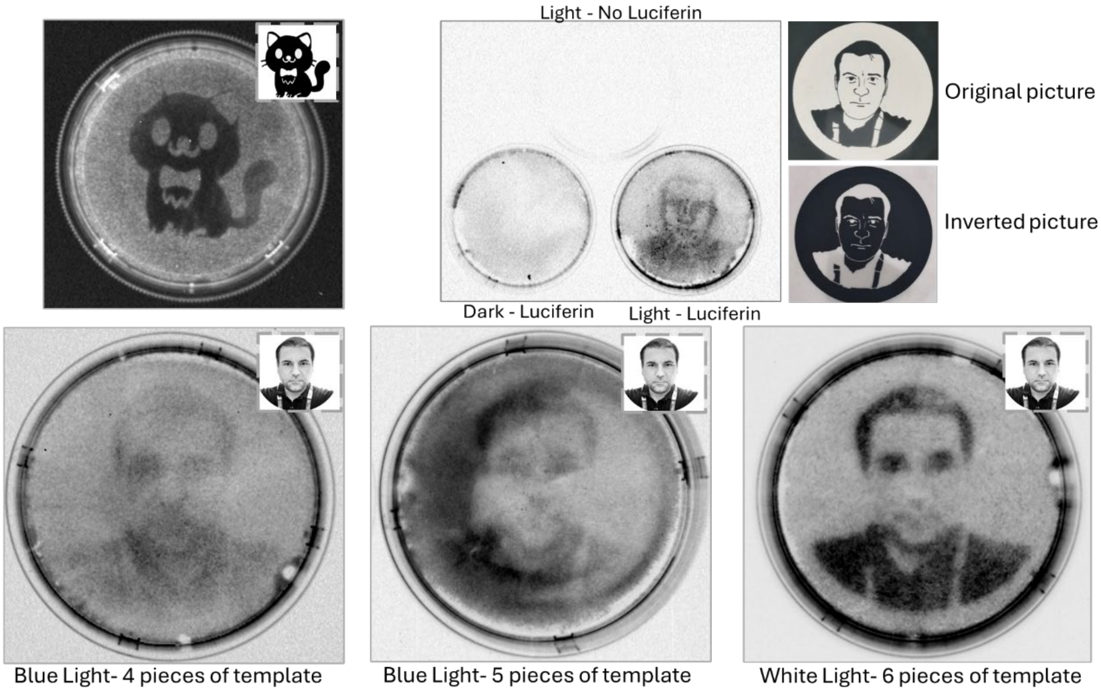


**Fig. S3**. Photomasks and imaging setups used to spatially control light exposure for pattern formation in cell culture. (*A*) Polystyrene photomasks were fabricated by cutting out the letters “CLK” and painting the remaining areas black to block light scattering. The panel shows an example of a “CLK” photomask placed on a 100-mm cell culture dish or “C”, “L”, and “K” photomasks placed on 35-mm cell culture dishes. Similar photomasks were prepared in different sizes to fit 100-mm and 35-mm dishes. Polystyrene photomasks were primarily used for initial optimization of pattern formation. (*B*) Top panel: Representative fluorescence images of cells transfected with cc-pcDNA4.TO-GFP and illuminated through the photomasks. Non-transfected cells were also exposed to light to confirm that no non-specific or autofluorescence signals were induced by white-light exposure. Transfected cells displayed general background fluorescence, but distinct patterns appeared only in light-treated samples. Bottom Panel: Similarly, cc-pcDNA4.TO-Luc–transfected cells were imaged for luminescence using the same imaging system employed for immunoblot chemiluminescence. All images were saved in grayscale, and signals were inverted for clarity and comparison between fluorescence and luminescence-based images. (*C*) For sketched images and human portraits, transparent acetate paper sheets were used as photomasks. The original images were inverted according to their black-white density and printed on clear acetate paper sheets; up to five layers were stacked to obtain the final photomask. The lower panel shows the original (non-inverted) forms of the images. (*D*) Example of raw, unprocessed images captured simultaneously to demonstrate the authenticity of pattern formation. Cropped (using crop to shape option) versions of these images are shown in Fig. 3 of the main text. Because the imaging field accommodated only one 100-mm dish during fluorescence imaging or two 100-mm dishes during luminescence imaging, images were captured sequentially but under identical acquisition settings. Partial images showing three dishes (fluorescence) or two full dishes (luminescence) were also recorded for visual comparison of dark and light samples. (*E*) To rule out any direct stimulatory effect of light on luciferase activity, cells were transfected with either a constitutively active plasmid (pcDNA4.TO-Luc) or a light-responsive plasmid (cc-pcDNA4.TO-Luc). Both dishes were exposed to the same “L”-shaped photomask and light source. As expected, the constitutive construct produced strong overall luminescence, whereas the light-responsive construct generated a distinct “L” pattern only upon illumination. Luminescence images were compared at exposure times (13.5 s vs. 300 s), yielding comparable background intensities. (*F*) Additional examples of luminescence-based patterning using acetate-sheet photomasks depicting a sketched cat and a human portrait which were generated with different photomasks than used in the main text and cells were illuminated using blue light instead of white light. To enhance image contrast prior to achieving realistic portraits, sketched versions derived from real human images were first employed; one such example is shown here. (*G*) Improvisation of human portrait formation in dishes. First two images were generated after being exposed to blue light, and images were generated by using 4 and 5 sheets of acetate paper while third image was generated by exposing cells to white light through 6 sheets of acetate paper. The best quality images taken using 5 sheets of acetate paper under white light were shown in the main manuscript file, Fig. 3*F*.

Figure S4

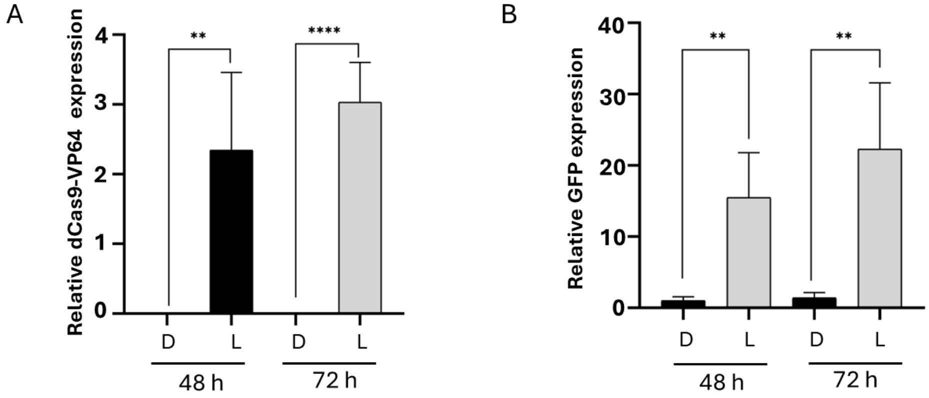


**Fig. S4.** Quantification and statistical analysis of CRISPR-based gene expression regulation by light. (*A*) Relative dCas9-VP64 expression levels in dark (D) and light (L) samples were quantified. (*B*) Relative GFP expression levels between D and L samples were also quantified. Statistical analyses for both dCas9-VP64 and GFP expression were performed using a two-tailed unpaired Student's *t*-test in GraphPad Prism. Results are presented as mean ± SD (n = 4). The levels of statistical significance were defined as: ns<0.12, *p<0,033, **p<0,002, ***p<0,001. The representative image given in Fig. 4*A* of the main manuscript file was one of the images which were used for the quantitation shown in this figure.

Figure S5

A

**Spectral properties of Light Source used for light treatments of 96- or 24-well plates and 35-mm dishes**

**Light Source Brand:** TaoTronics **Model:** Dimmable Touch Eye-protection Led Desk Lamp-TT-DL13

**Light meter Brand**: AquaHorti **Model**: AH-300 Light Meter.

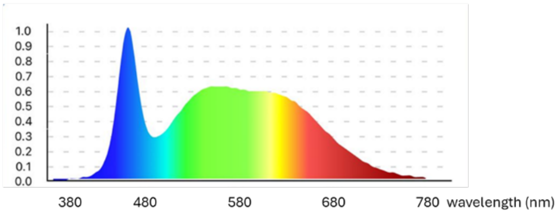


B **Spectral Properties of light source used for image formation studies in 60-mm or 100-mm culture dishes**

**Light Source Brand:** GQP **UPC:**683404119938 **Model:** Light Therapy Lamp (UV-Free Sun Lamp with 660nm Red Light and 12000 Lux Bright White Light)

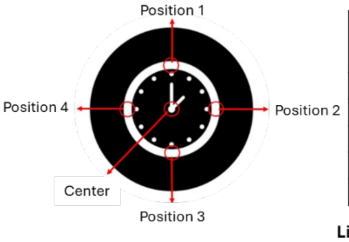


| Position | Target (lux) | Mean (lux) | SD (lux) | Converted_Irradiance_ W/m² | J/m²_24h | Temperature |
|---|---|---|---|---|---|---|
| 1 | 1000 | 964,7 | 6,429 | 2,923 | 252576,000 | 35,04 |
| 2 | 1000 | 954,7 | 5,033 | 2,893 | 249957,818 | 35,01 |
| 3 | 1000 | 985 | 12,17 | 2,985 | 257890,909 | 35 |
| 4 | 1000 | 1007 | 4,583 | 3,052 | 263650,909 | 35,02 |
| Center | 1000 | 1016 | 4,163 | 3,079 | 266007,273 | 35 |

**Light meter Brand:** Fisher Scientific **Model:** Fisherbrand™ Traceable™ Dual-Display Light Meter reader

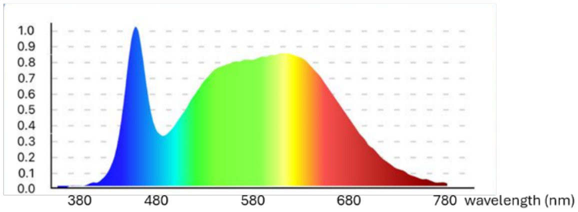


**Fig. S5.** Properties of light sources and their spectral profiles. (*A*) Cells cultured in 96-well, 24-well, or 35-mm cell culture dishes were illuminated using a reading desk LED lamp. Its spectral profile is shown. (*B*) Because the desk lamp did not uniformly illuminate cell culture dishes larger than 35-mm, a table-shaped LED source was used for larger culture areas. Light intensity was measured at multiple positions under the photomask to confirm homogeneity, and a representative emission spectrum is shown.
